# *In vivo* and *in vitro* mechanistic characterization of a clinically relevant PolγA mutation

**DOI:** 10.1101/2020.09.10.291369

**Authors:** Pedro Silva-Pinheiro, Carlos Pardo-Hernández, Aurelio Reyes, Lisa Tilokani, Anup Mishra, Raffaele Cerutti, Shuaifeng Li, Dieu Hien Ho, Sebastian Valenzuela, Anil Sukru Dogan, Peter Bradley, Patricio Fernandez-Silva, Aleksandra Trifunovic, Julien Prudent, Michal Minczuk, Laurence Bindoff, Bertil Macao, Massimo Zeviani, Maria Falkenberg, Carlo Viscomi

## Abstract

Mutations in *POLG*, encoding POLγA, the catalytic subunit of the mitochondrial DNA polymerase, cause a spectrum of disorders characterized by mtDNA instability. However, the molecular pathogenesis of *POLG*-related diseases is poorly understood and efficient treatments are missing. Here, we generated a *POLG^A449T/A449T^* mouse model, which reproduces the most common human recessive mutation of *POLG*, encoding the A467T change, and dissected the mechanisms underlying pathogenicity. We show that the A449T mutation impairs DNA binding and mtDNA synthesis activities of POLγ *in vivo* and *in vitro*. Interestingly, the A467T mutation also strongly impairs interactions with POLγB, the homodimeric accessory subunit of holo-POLγ. This allows the free POLγA to become a substrate for LONP1 protease degradation, leading to dramatically reduced levels of POLγA, which in turn exacerbates the molecular phenotypes of *Polg^A449T/A449T^* mice. Importantly, we validated this mechanism for other mutations affecting the interaction between the two POLγ subunits. We suggest that LONP1 dependent degradation of POLγA can be exploited as a target for the development of future therapies.

## Introduction

*POLG*, encoding the catalytic subunit of mitochondrial DNA-specific polymerase gamma (POLγA), contains the highest number of deleterious mutations of the human coding genome, associated with a huge spectrum of clinical, molecular and biochemical phenotypes. Autosomal dominant and recessive mutations have been reported, and among the recessive ones, by far the most common is a missense mutation changing A467 into a T, in an area of the protein which has no obvious function. Nevertheless, the consequences of this mutation are devastating in humans, including severe, juvenile neurodegeneration of the spinal cord, brainstem, neocortex, particularly in the occipital pole, associated with ataxia, refractory epilepsy, toxicity to valproate, cognitive regression and eventually global neurological impairment and death. Status epilepticus is a common end conclusive outcome of the homozygous patients. The A467T is the most frequent POLγA mutation in Scandinavian and Northern European Countries, together with another change, the W748S change, which seems to be part of the Finnish disease heritage. In both cases, no obvious alteration of relevant known function of the primary structure of the protein seems to be affected, neither the proofreading activity present in the N-terminus of the protein, downstream from the mitochondrial targeting sequencing, nor the polymerase domain, which is confined to the C-terminus of the sequence. The intermediate region, where the two deleterious mutations are located, is generically defined as the “linker” region between the proofreading and the polymerase domain, and seems to play a role, not better specified, in binding the second component of POLγ, the 55 kDa subunit PolγB, which binds homodimerically POLγA, conferring to the whole trimeric mtDNA replisome a marked increase in polymerase processivity. Therefore, the mechanistic abnormalities by the two most common recessive mutations affecting the activity of this important enzyme are virtually unknown. Notably, the A467T mutation, when associated with an allelic null mutation leads to an even more devastating condition named Alpers-Huttenlocher syndrome, which affects babies suffering of refractory epilepsy due to severe spongiotic atrophy of the brain, and hepatic failure. As already mentioned, in addition to the A467T and W748S, over 300 mutations have been described in *POLG* (Human DNA Polymerase Gamma Mutation Database: https://tools.niehs.nih.gov/polg/) leading to a spectrum of diseases, which has been tentatively classified in the following categories: (i) the already mentioned Alpers-Huttenlocher syndrome (AHS), (ii) myocerebrohepatopathy spectrum (MCHS), which presents with developmental delay, lactic acidosis, myopathy and hepatic impairment; (iii) the typical spectrum of the A467T and W748S mutations (in either homozygosity or in combination), including myoclonic epilepsy myopathy sensory ataxia (MEMSA), comprising spinocerebellar ataxia with epilepsy (SCAE), frequently associated with sensory ataxia neuropathy with dysarthria and ophthalmoplegia (SANDO), and, (iv) finally, autosomal dominant and recessive progressive external ophthalmoplegia (ad and arPEO) (Rahman & Copeland, 2019). The plethora of terms defining the different conditions of POLγA mutations reflects the huge spectrum of syndromic presentations associated with abnormalities of this essential enzyme. Consequently, it is difficult to rigidly classify these syndromes as often symptoms overlap, the same mutation can be associated to more than one presentation, and each phenotype can be the consequence of the combination of different allelic mutations. In several cases, particularly in syndromes associated with progressive external ophthalmoplegia, abnormalities of the mtDNA integrity lead to the accumulation of multiple mtDNA deleted species; in other cases, such as the A467T mutation itself, these large scale rearrangements are rare, but whenever necropsy examination has been performed, depletion of mtDNA was found in critical areas, particularly in specific regions of the brain (Tzoulis, Tran et al., 2014). Again, the mechanistic details leading to the generation of these molecular lesions is poorly understood.

As already mentioned, DNA polymerase γ (POLγ) is a heterotrimer with one catalytic POLγA subunit and two POLγB accessory subunits (Gustafsson, Falkenberg et al., 2016). The *POLG* gene codes for the 140 kDa POLγA subunit that harbors DNA polymerase, 3′-5′ exonuclease, and 5′-deoxyribose phosphate lyase activities (Longley, Prasad et al., 1998), whereas *POLG2* encodes the 55 kDa POLγB, which stabilizes the interactions with template-DNA, thereby increasing processivity (Lim, Longley et al., 1999). POLγ is the only DNA polymerase required for mtDNA replication in mammalian mitochondria and, at the replication fork, it works in concert with the TWINKLE DNA helicase (Spelbrink, Li et al., 2001). The mitochondrial single-stranded DNA-binding protein (mtSSB) stimulates mtDNA synthesis by increasing the helicase activity of TWINKLE and the DNA synthesis activity of POLγ (Korhonen, Gaspari et al., 2003).

Four mutations (A467T, W748S, G848S and the T251I–P587L allelic pair) account for ~50% of all mutations identified in patients with *POLG*-related diseases, with ~75% of patients carrying at least one of these mutant alleles (Uusimaa, Gowda et al., 2013). *POLG* mutations may lead to mtDNA instability, causing either multiple deletions or depletion ^2^. However, there is no obvious genotype– phenotype correlation, and, as mentioned above, the same mutation can often lead to mtDNA deletions, mtDNA depletion or both. A prototypical example is highlighted by the homozygous mutation A467T mutation, which has been associated with a range of phenotypes, from childhood-onset fatal AHS to MEMSA, ANS and SANDO (Rahman & Copeland, 2019). In addition, the age of onset and the progression of POLG-related disease in patients with the same *POLG* mutations is astonishingly variable and can span several decades. For instance, the onset of disease spans >70 years in compound heterozygous patients carrying the T251I–P587L mutations on one allele (DeBalsi, Longley et al., 2017) and the G848S mutation on the other, and it spans at least four decades of life in homozygous A467T patients (Rajakulendran, Pitceathly et al., 2016, Tzoulis, Engelsen et al., 2006). The A467T is the most common pathogenic variant of *POLG* (de Vries, Rodenburg et al., 2007, Ferrari, Lamantea et al., 2005, Horvath, Hudson et al., 2006, Nguyen, Sharief et al., 2006), and it seems to severely impair POLγ activity (Chan, Longley et al., 2005). The proposed, but not firmly established, mechanisms include (i) reduction of the affinity of the enzyme for deoxynucleotide triphosphates (dNTPs), (ii) lowering of the catalytic activity, and (iii) reduction of the affinity for the *POLG2*-encoding accessory subunit. All this would lead to stalling of the replication fork and depletion or instability of mtDNA. Nevertheless, mechanistic evidence of these effect remains largely hypothetical. It is therefore crucial for the understanding of the pathogenic mechanisms encompassing this huge area of neurodegenerative disorders, and the dissection of the very function of several structures and interactions of POLγA, to better understand the mechanistic processes leading to POLγ impairment in the presence of a change that seems not to compromise known essential function of the protein, but that nevertheless retains an obviously relevant physiological function and has an essential impact in the medicine associated with POLγ impairment.

Here, we generated a *Polg^A449T/A449T^* knockin mouse by CRISPR/Cas9 technology, which reproduces the A467T human mutation. By using a combination of cellular, *in vivo* and *in vitro* techniques, we show that the mutation not only impacts on polymerase activity, DNA binding and interaction with PolγB, but also makes the protein exquisitely susceptible to degradation by the LONP1 protease, the most important soluble protease of the mitochondrial matrix. Our results thus reveal a novel pathogenic mechanism for *POLG*-related diseases, which in turn suggests new avenues for development of future therapies.

## Results

### Generation and characterization of *Polg^A449T/A449T^* mutant mice

To investigate the molecular pathogenesis of *POLG*-related disorders, we generated a *Polg^A449T/A449T^* homozygous knockin mouse, corresponding to the human A467T mutation, by CRISPR/Cas9 technology (Supplementary Figure 1). Three-month old *Polg^A449T/A449T^* homozygous animals did not show any gross phenotype compared to wild-type (WT) littermates, including similar body weight curve and rotarod performance (not shown). However, a significant reduction in treadmill motor endurance was detected (Figure 1A). Although whole body metabolism was similar in KI and controls by CLAMS analysis, a significant reduction in spontaneous rearing movements was observed in *Polg^A449T/A449T^* mutants (Figure 1B-C and Supplementary Figure 2). Post-mortem hematoxylin and eosin staining did not show any gross abnormality in any tissue.

**Figure 1.**
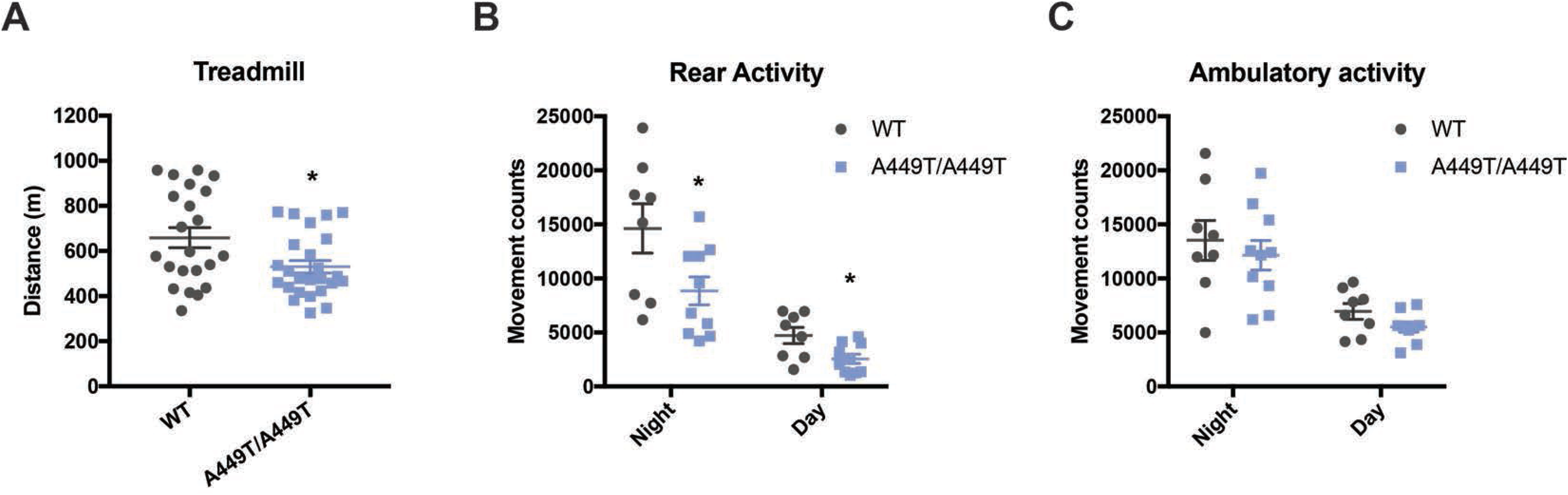
Characterization of the clinical phenotype of *Polg^A449T/A449T^* mice. **A.** Distance run in metres by 3 month-old WT and *Polg^A449T/A449T^* animals on the treadmill. Data are presented as mean ± SEM. *p<0.05; Student’s t-test. Each symbol represents a biological replicate. **B.** Spontaneous rear activity (vertical movement counts) of 3 month-old WT and *Polg^A449T/A449T^* animals measured in the CLAMS™ system. Data are presented as mean ± SEM. *p<0.05; Student’s *t*-test. Each symbol represents a biological replicate. **C.** Spontaneous ambulatory activity (horizontal movement counts) of 3 month-old WT and *Polg^A449T/A449T^* animals measured in the CLAMS™ system. Data are presented as mean ± SEM. Student’s *t*-test. Each symbol represents a biological replicate.

**Figure 2.**
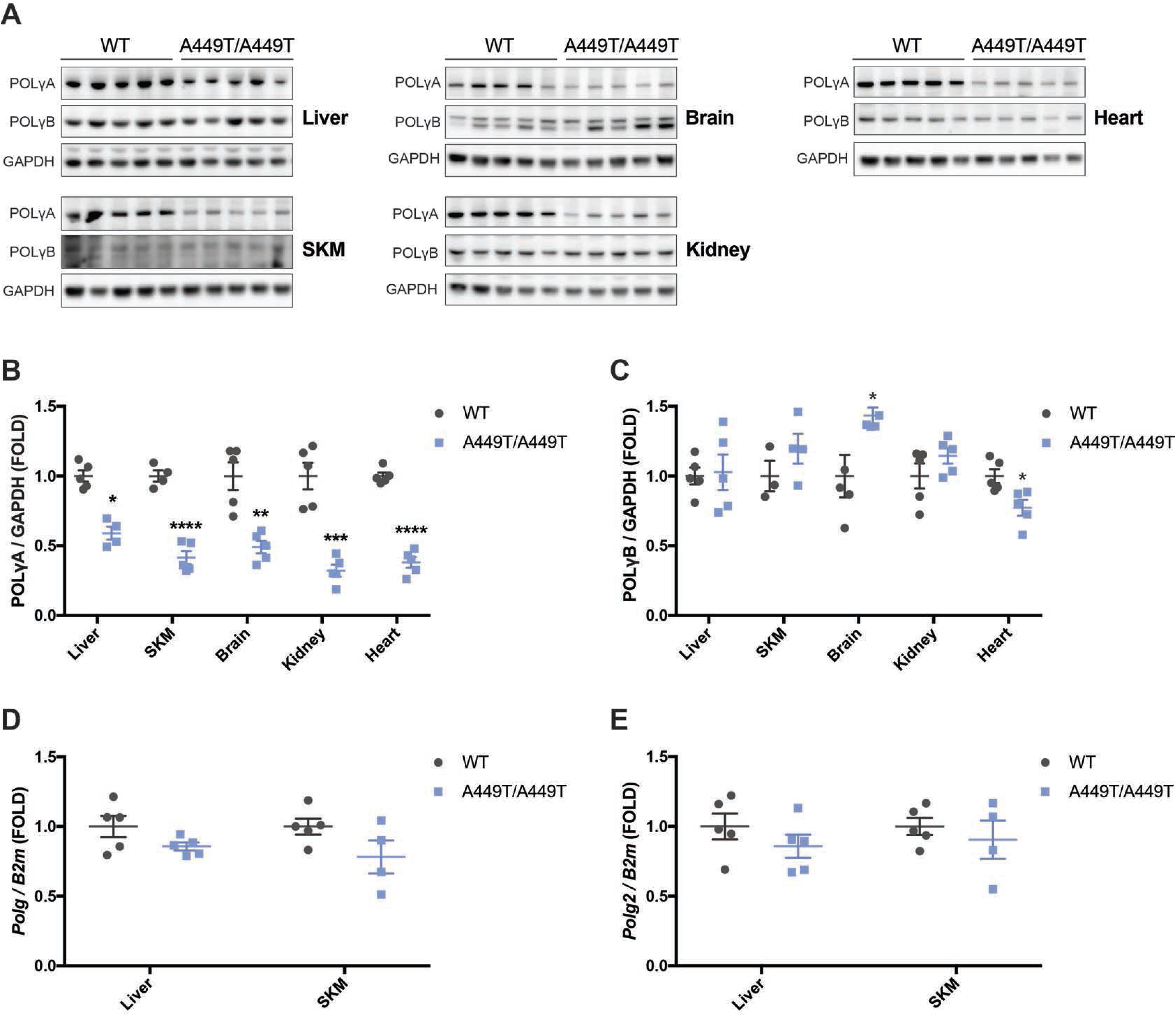
Characterization of PolγA and PolγB levels in tissues of *Polg^A449T/A449T^* mice. **A.** Western blot analysis of steady-state levels of PolγA and PolγB in liver, skeletal muscle (SKM), kidney, brain and heart of WT and *Polg^A449T/A449T^* animals. The lower band in the brain is an aspecific. GAPDH was used as loading control. **B-C.** Quantification of (**A**). PolγA (**B**) and PolγB (**C**) levels were normalized to GAPDH and presented as FOLD change from WT animals. Data are presented as mean ± SEM. *p<0.05; **p<0.01; ***p<0.001; ****p<0.0001; Student’s *t*-test. Each symbol represents a biological replicate. **D-E.** Real-Time qRT-PCR quantification of the transcripts *Polg* **(D)**and *Polg2* **(E)**, normalized to *B2m*, in liver and skeletal muscle (SKM) of WT and *Polg^A449T/A449T^* animals. Data are presented as mean ± SEM. Student’s *t*-test. Each symbol represents a biological replicate.

### POLγA is reduced in*Polg^A449T/A449T^* tissues

We analysed the effects of the A449T mutation on POLγA and POLγB protein levels. Immunoblotting revealed strong reduction of PolγA^A449T^ amount, as low as 50%, in all tissues examined, including brain, heart, skeletal muscle, liver and kidney (Figure 2A-B). In contrast, POLγB levels were unchanged in most tissues, albeit a mild upregulation and downregulation in brain and heart, respectively, was observed (Figure 2A-C). Analysis of the corresponding mRNAs showed no significant changes of *Polg* or *Polg2* transcripts in both liver and skeletal muscle of *Polg^A449T/A449T^* compared to control littermates (Figure 2D-E), suggesting post-translational instability of the mutant protein.

### Reduced mtDNA content and impaired replication in *Polg^A449T/A449T^* tissues and MEFs

Since mutations in *POLG* are associated with mtDNA instability in human patients, we next investigated mtDNA content and integrity in several tissues, including liver, skeletal muscle, brain, kidney and heart from both *Polg^A449T/A449T^* vs. WT littermates (Figure 3A). MtDNA copy number was significantly reduced in the skeletal muscle of *Polg^A449T/A449T^* (80±4%, p˂0.01) compared to WT littermates, without accumulation of multiple deletions (Figure 3B and Supplementary Figure 3A-C). No difference in mitochondrial transcripts and OXPHOS activities were detected between *Polg^A449T/A449T^* vs. WT littermates in liver and SKM (Supplementary Figure 3D-J).

**Figure 3.**
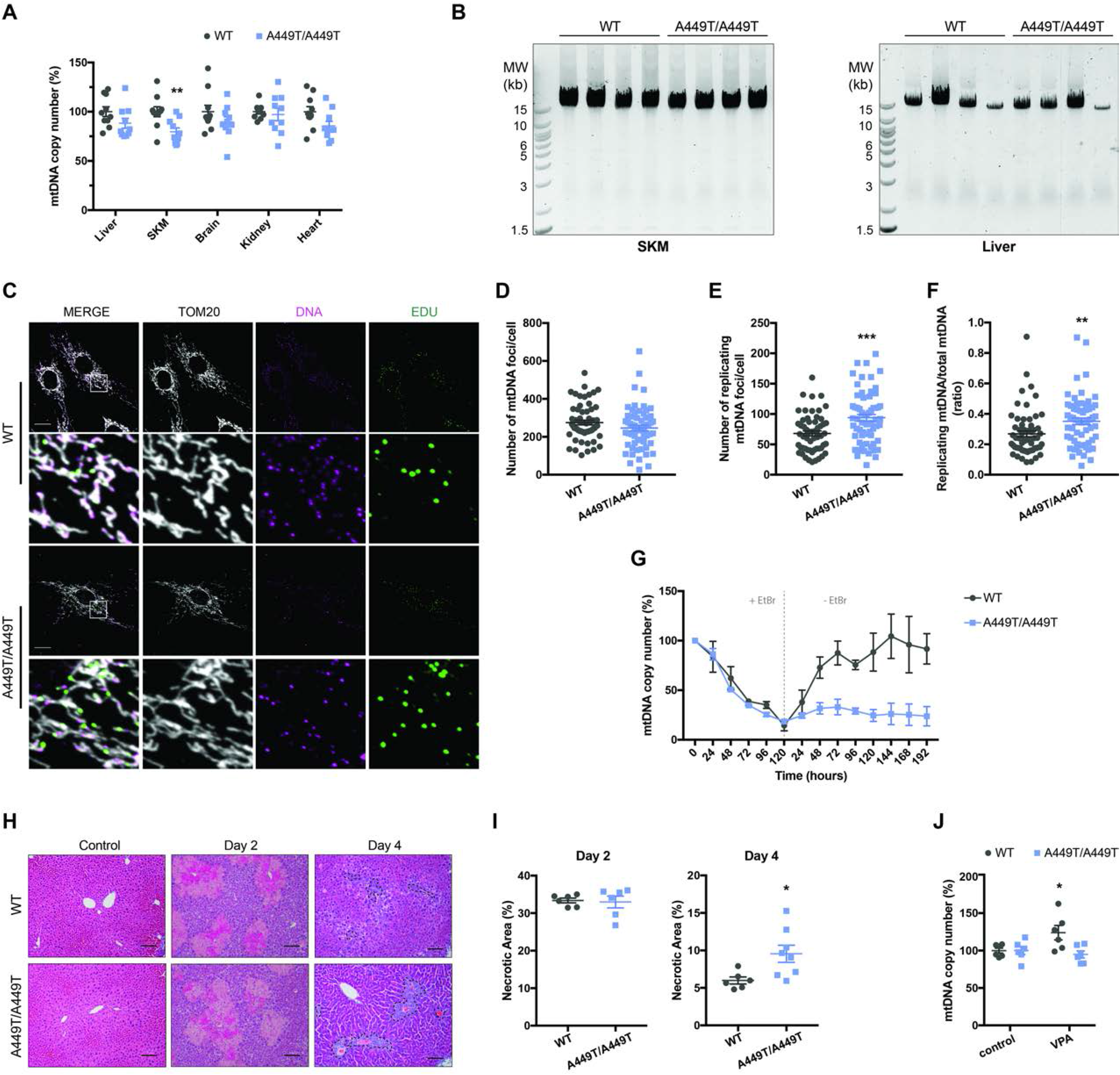
Characterization of the molecular phenotype of *Polg^A449T/A449T^* mice. **A.** Real-Time qPCR quantification of mtDNA content in liver, skeletal muscle (SKM), kidney, brain and heart of WT and *Polg^A449T/A449T^* animals. Data are presented as mean ± SEM. **p<0.01; Student’s *t*-test. Each symbol represents a biological replicate. **B.** Long-range PCR performed in DNA isolated from skeletal muscle (SKM) and liver of WT and *Polg^A449T/A449T^* animals. Primers amplifying a fragment of 15,781bp of the mtDNA. The bands were visualized by SYBR™ safe staining. **C.** Representative confocal images of mitochondria, DNA and replicating mtDNA (EdU) from WT and *Polg^A449T/A449T^* MEFs. Mitochondria and mtDNA were labelled using anti-TOM20 and anti-DNA antibodies, respectively. Replicating DNA was visualized in fixed cells after incubation with 50 μM EdU for 1 hour. Scale bar 20 μm. **D.** Quantification of total mtDNA from (**A**). Data are presented as mean ± SEM. Student’s *t*-test. Each symbol represents individual cells (n=60) from three independent experiments. **E.** Quantification of mitochondrial EdU positive foci from (**A**). Data are presented as mean ± SEM. ***p<0.001; Student’s *t*-test. Each symbol represents individual cells (n=60) from three independent experiments. **F.** Ratio of the mitochondrial replicating mtDNA / total mtDNA. Data are presented as mean ± SEM. **p<0.01; Student’s *t*-test. Each symbol represents individual cells (n=60) from three independent experiments. **G.** Real-Time qPCR quantification of mtDNA content in WT and *Polg^A449T/A449T^* MEFs during Ethidium Bromide-mediated depletion and then recovery of mtDNA. Data are presented as mtDNA percentage (%) of untreated cells of each genotype. Data are presented as mean ± SEM. (n = 3). **H.** Representative H&E staining of liver tissue sections of WT (top) and *Polg^A449T/A449T^* (bottom) animals with a single injection of CCl_4_. Two days after injection (middle), 4 days after injection (right) and control/non-injected mice (left). Note the necrotic areas around the central veins. Scale bar 100 μm. **I** Quantification of necrotic areas (D) as percentage (%) of the total section area, 2 and 4 days after a single injection of CCl_4_. Data are presented as mean ± SEM. *p<0.05; Student’s t-test. Each symbol represents a biological replicate. **J.** Real-Time qPCR quantification of mtDNA content in liver of control and VPA-treated WT and *Polg^A449T/A449T^* mice for 60 days. Data are presented as mean ± SEM. *p<0.05; Student’s *t*-test. Each symbol represents a biological replicate.

To investigate in detail the effects on mtDNA replication we generated mouse embryonic fibroblasts (MEFs) from *Polg^A449T/A449T^* and WT cells. The mtDNA content was similar in the two genotypes (Supplementary Figure 3K). We then investigated mtDNA replication in MEFs, using 5-ethynyl-2′-deoxyuridine (EdU) staining in junction with an anti-DNA antibody to label replicating and total mtDNA. Interestingly, we observed a significantly increased fraction of replicating mtDNA molecules in *Polg^A449T/A449T^* vs. WT MEFs (Figure 3C-F), indicating that more mtDNA foci were engaged in replication. Next, we used ethidium bromide (EtBr) to deplete mtDNA content, and found that after removal of EtBr, mtDNA content recovered to pre-treatment values within 3 days in WT MEFs, whereas no recovery at all was observed in the mutant cells (Figure 3G), strongly indicating severely impaired mtDNA replication in stress conditions of *Polg^A449T/A449T^* mouse mitochondria.

Given the mild reduction in mtDNA copy number in the mutant mice, we decided to challenge them with a single injection of carbon tetrachloride (CCl_4_), which induces acute liver damage, triggering liver cell division to repopulate the necrotic areas. Two days after the injection, both WT and *Polg^A449T/A449T^* showed extensive areas of necrosis (approximatively 35% of the liver), which was reduced to 6±0.46 % in WT mice after four days, whereas it was still above 10% in *Polg^A449T/A449T^* mice (10±1.15%, p˂0.05) (Figure 3H-I). This result clearly indicates that cell replication is impaired in *Polg^A449T/A449T^*, likely due to lack of bioenergetic supply by impaired mitochondria in stress conditions. Since the antiepileptic drug valproic acid (VPA) is known to induce acute liver failure in patients with the A467T mutation in POLγA (Saneto, Lee et al., 2010, Stewart, Horvath et al., 2010), we treated our mice with VPA by daily oral gavage (300 mg/kg) for one week or in food pellets (1.5% VPA) for two month. Both WT and *Polg^A449T/A449T^* did not show any sign of hepatic failure or histological damage. However, whilst WT-treated mice showed increased mtDNA copy number compared to non-treated WT mice (124±9.7%, p˂0.05) after chronic treatment, the mtDNA content in *Polg^A449T/A449T^*-treated mice was similar to non-treated mutants (94.9±4.7%, p=0.4889), suggesting reduced mtDNA replication in response to VPA challenge (Figure 3J).

### Polg^*A449T/A449T*^ mitochondria have reduced 7S DNA and accumulate replication intermediates

We then investigated mtDNA replication in the tissues of the mutant and control mice by Southern blot. Normally, about 95% of all replication events are prematurely terminated, generating a 650 nucleotide-long molecule, called 7S DNA (Bogenhagen & Clayton, 1978, Doda, Wright et al., 1981, Nicholls & Minczuk, 2014). In *Polg^A449T/A449T^* but not in WT littermate, the 7S DNA levels were significantly reduced in skeletal muscle and kidney, and a similar trend was also present in the other analysed tissues, except for the heart, (Figure 4A-B and Supplementary Figure 4A-C). These results suggest compensatory mtDNA replication in knockin mice vs. WT littermates.

**Figure 4.**
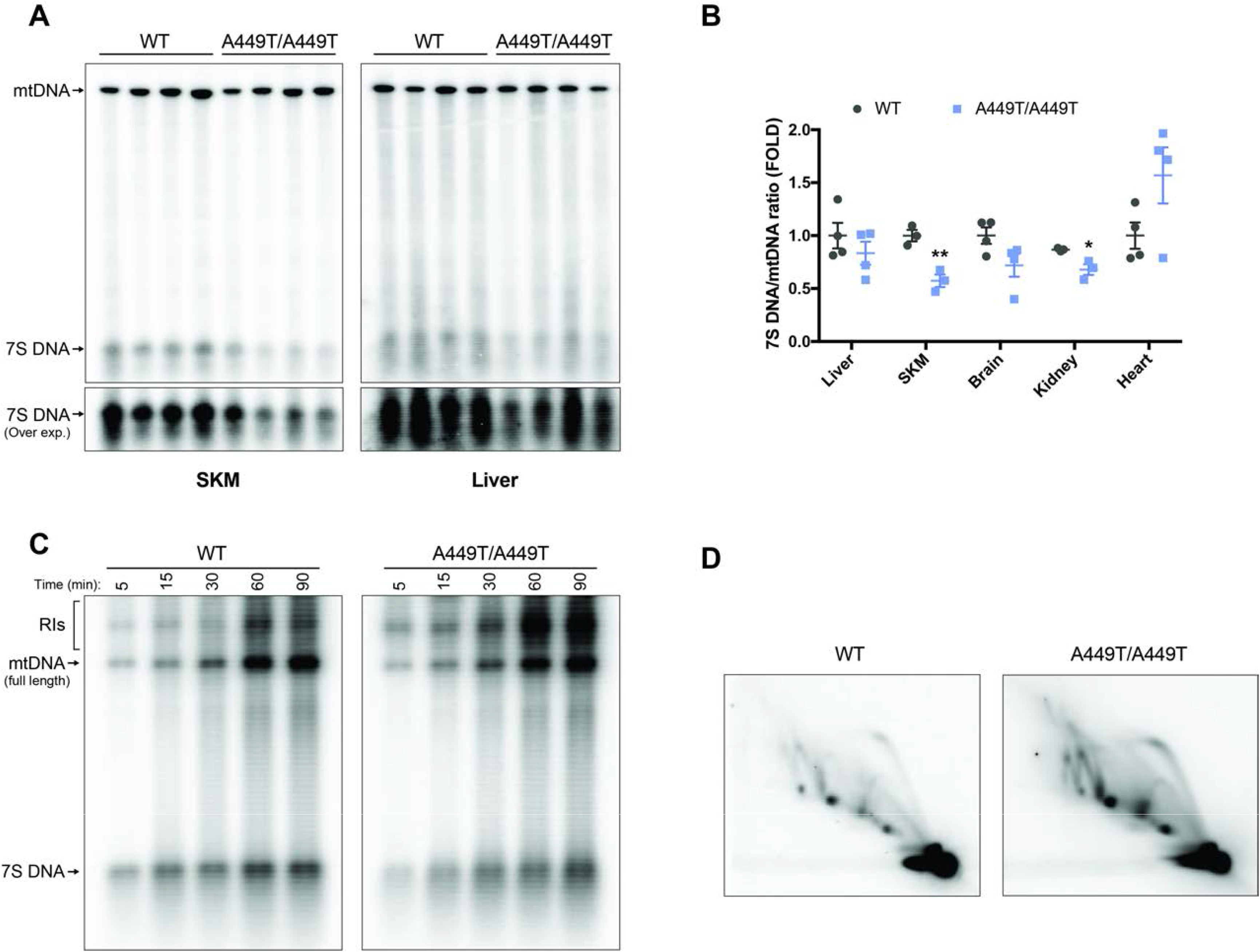
Molecular analysis of mtDNA replication in *Polg^A449T/A449T^* mitochondria. **A.** Southern blot analysis of BlpI-digested mtDNA and 7S DNA from skeletal muscle (SKM) and liver of WT and *Polg^A449T/A449T^* animals. **B.** Quantification of the Southern blots presented in panel A and Supplemental Figure 4A-C. 7S DNA levels were normalized to linearized full length mtDNA and presented as FOLD change from WT animals. Data are presented as mean ± SEM. *p<0.05; **p<0.01; Student’s *t*-test. Each symbol represents a biological replicate. **C.** Time course of *de novo* DNA synthesis of mtDNA, 7S DNA and RIs in liver-isolated mitochondria of WT and *Polg^A449T/A449T^* animals. Brackets indicate mtDNA replication intermediates (RIs). Pulse-labelling time (minutes) is indicated on the top. **D.** Analysis of the mtDNA replication intermediates (RIs) in the liver of WT and *Polg^A449T/A449T^* mice, resolved by 2D-AGE and followed by southern blot visualization. DNA was digested with the BclI restriction enzyme. For probe and restrictions sites location, schematic representation and quantification of the different types of RIs, please refer to (Supplemental Figure 4D-F).

To better investigate the mechanistic details of mtDNA replication, we then performed *in organello* replication experiments in isolated liver mitochondria (Figure 4C), by pulse-labelling with α-^32^P-dATP. Although no obvious differences were detected in mtDNA replication rates between *Polg^A449T/A449T^* and WT mice (Figure 4C), the signal due to long but incomplete mtDNA molecules was much more intense in the *Polg* mutant compared to WT samples, thus suggesting accumulation of replication intermediates (RIs) in the mutant vs. controls. Accordingly, we applied two-dimension agarose gel electrophoresis (2D-AGE), which resolves DNA molecules based on size and shape, allowing a snapshot of the RIs. Notably, *Polg^A449T/A449T^* mice displayed an overall accumulation of the different types of RIs compared to WT animals (Figure 4D and Supplementary Figure 4D-F), revealing abnormal replication of *Polg^A449T/A449T^* mainly due to generalized replication fork stalling.

These results are concordant with those found in MEFs (Figure 3C-F).

These novel data clearly demonstrate that the A449T mutation impairs mtDNA replication in both cultured cells and in vivo.

### Polγ^AA449T^ protein has reduced affinity for DNA which is partially rescued by PolγB subunit

To further document the stalling phenotype of the A449T mutant in vitro, we expressed and purified WT and mutant PolγA as recombinant proteins. First, we used an electrophoretic mobility shift assay (EMSA) to measure the binding of Polγ to a primed DNA template. When alone, POLγA^A449T^ bound DNA ≈ 3.4 times more weakly than POLγA^WT^ (Figure 5A and Supplemantary Figure 5A) and remained substantially lower than the WT also after the addition of POLγB (Figure 5B and Supplementary Figure 5B).

**Figure 5.**
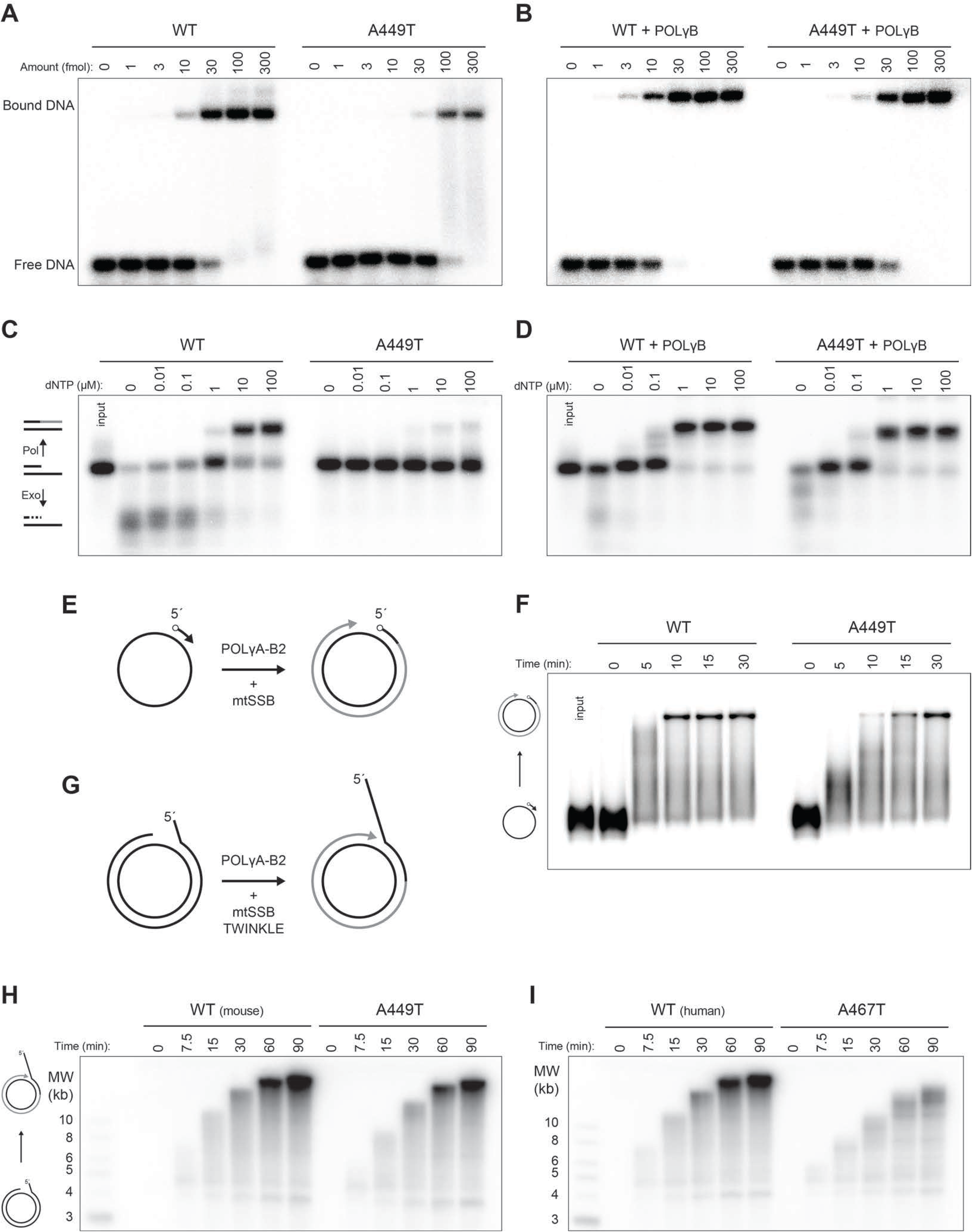
*In vitro* characterization of POLγA^A449T^ mutant protein. **A.** Electrophoretic mobility assays using POLγA^WT^ and mutant PolγA^A449T^ to estimate affinity to a DNA template. For Kd (DNA) calculations please refer to (Supplemental Figure 5A). Each lane contains 10 fmol of DNA substrate and the indicated amounts of PolγA on the top. **B.** Electrophoretic mobility assays using POLγA^WT^ and mutant POLγA^A449T^ with addition of POLγB to estimate affinity to a DNA template. For Kd(DNA) calculations please refer to (Supplemental Figure 5B). Each lane contains 10 fmol of DNA substrate and the indicated amounts of POLγ holoenzyme on the top. **C.** Coupled exonuclease-polymerase assay using POLγA^WT^and mutant POLγA^A449T^ across increasing concentrations of dNTPs using a short DNA template. A schematic representation of the assay is presented on the left. **D.** Coupled exonuclease-polymerase assay using POLγA^WT^ and mutant POLγA^A449T^ with addition of POLγB, across increasing concentrations of dNTPs using a short DNA template. **E.** Schematic representation of the second strand synthesis assay. This assay evaluates the ability of polymerise long stretches of DNA by synthesising the second strand of a single stranded template hybridized with a 5’ radiolabelled primer. MtSSB is added in the reaction. **F.** Second strand synthesis assay using WT and mutant POLγA^A449T^ to assess polymerase activity using longer DNA templates. The reactions include POLγA-B2 and mtSSB and were incubated for the indicated times on top of the blot. **G.** Schematic representation of the Rolling circle *in vitro* replication assay. The template consists of an incomplete double stranded DNA template with a mismatch on the 5’ of the incomplete strand. In the presence of TWINKLE and mtSSB, POLγA-B2 can polymerase long stretches of DNA using the 3’-end of the incomplete strand. **H.** Rolling circle *in vitro* replication assay using POLγA^WT^ and mutant POLγA^A449T^ to assess polymerase activity in the context of the minimal mitochondrial replisome, which includes POLγ holoenzyme WT or mutant, TWINKLE and mtSSB). The reactions were incubated for the indicated times (top). **I.** Rolling circle *in vitro* replication assay using human versions of POLγA^WT^ and mutant POLγA^A467T^ to assess polymerase activity in the context of the minimal mitochondrial replisome (POLγ holoenzyme WT or mutant, TWINKLE and mtSSB). The reactions were incubated for the indicated times (top).

### POLγ^AA449T^ has reduced polymerase activity which is partially rescued by POLγB

Next, we investigated POLγA activities using a short DNA template annealed to a radioactively labelled primer. By performing the experiment across a range of dNTP concentrations, we could analyze both polymerase and exonuclease function. The exonuclease activity can digest the labelled primer, whereas the polymerase activity can elongate the primer and synthesize an additional short, 15 nucleotide stretch of DNA. As expected, at lower dNTP levels, POLγA^wt^ displayed 3’-5’ exonuclease activity, but at higher concentrations, it switched to polymerase activity (Figure 5C). Addition of POLγB reduced exonuclease activity and favored DNA synthesis even at lower dNTP concentrations (Figure 5D). The mutant POLγA^A449T^ was completely inactive in isolation, most likely due to its inability to efficiently bind primed DNA (Figure 5C). Nevertheless, addition of POLγB restored the polymerase activities of POLγA^A449T^, to levels similar to those observed with POLγA^wt^ (Figure 5D), whereas exonuclease activity was reduced also in POLγA^wt^ as a consequence of predominant polymerase activity measured in vitro (Figure 5D).

To further challenge the system, we performed a DNA synthesis assays using a long circular ssDNA template of 3000 nts (Figure 5E). POLγA^A449T^ displayed a clearly slower DNA synthesis rate compared to the POLγA^WT^, even in the presence of the POLγB subunit (Figure 5F). In order to monitor the effects of the A449T mutation on replication of dsDNA, we used a template containing a ~4 kb long dsDNA region with a free 3′-end acting as a primer (Figure 5G). Addition of the TWINKLE DNA helicase was required to unwind the DNA and the reaction was stimulated by mtSSB (Figure 5G). This reaction is absolutely dependent on POLγB and once initiated, very long stretches of DNA can be formed. In this rolling circle replication assay, POLγA^A449T^ showed reduced polymerase DNA synthesis rate (Figure 5H) compared to POLγA^WT^, at all concentrations tested (Supplementary Figure 5C), demonstrating that POLγA^A449T^ has reduced polymerase activity. This in vitro result is in perfect agreement with the stalling phenotype seen *in vivo*. A similar effect was obtained with the human POLγA^A467T^ (Figure 5I).

Analysis of incorporated radiolabelled nucleotides over time indicated that the *in vitro* replication rates with 10 μM dNTPs, were reduced to about 60% for POLγA^A449T^ compared to POLγA^WT^ (3.5 fmol/min vs 5.5 fmol/min) (Supplementary Figure 5D and 5E). Interestingly, the reduction was more pronounced with human POLγA^A467T^ compared to human POLγA^WT^ (1.4 fmol/min vs 5.3 fmol/min), than for the mouse equivalents, which could explain the more severe phenotype observed in patients (Supplementary Figure 5D and 5E).

### POLγA is unstable in absence of POLγB

The amount of POLγA^A449T^ was reduced in the *Polg^A449T/A449T^*, which could also contribute to impaired mtDNA replication. To better understand the impact of the A449T mutation on protein stability, we performed a thermofluor stability assay and monitored temperature-induced unfolding of POLγA^WT^ and POLγA^A449T^, both in absence and in presence of POLγB (Figure 6A and 6B). The stability assay revealed no major differences in the fluorescence profile between POLγA^WT^ and POLγA^A449T^ from 37°C upwards, but the fluorescence signal of POLγA^A449T^ was already higher than the WT at 25°C, clearly indicating that the mutant protein was already partially unfolded even at <37°C temperatures (Figure 6A and 6B). Interestingly, the presence of the POLγB had a dramatic stabilizing effect, by increasing the unfolding temperature of about 10 °C for both proteins (Figure 6A and 6B). These data suggest that also POLγA^WT^ is partially unstable in the absence of POLγB. Accordingly, a recent report demonstrated that human POLγB-knockout cells showed severe decrease in POLγA levels (Do, Matsuda et al., 2020).

**Figure 6.**
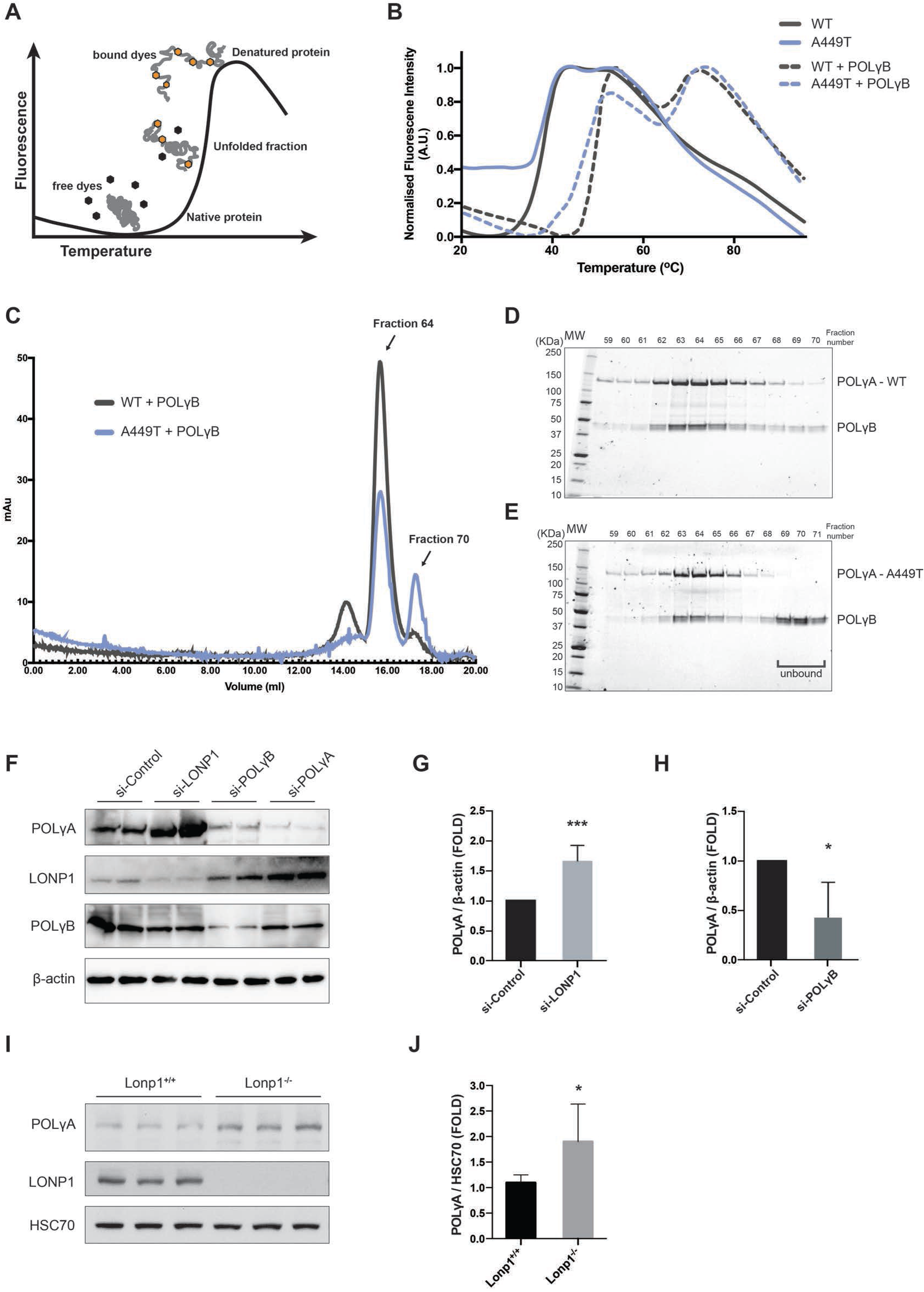
Stability of PolγA^A449T^ mutant protein *in vitro* and *in vivo*. **A.** Schematic representation of a typical thermofluor stability assay. This assay uses a fluorescent dye, SYPRO Orange, to monitor the temperature-induced unfolding of proteins. When the temperature starts to rise and unfold the protein, the SYPRO Orange dye fluoresces by binding to exposed hydrophobic patches. **B.** Thermofluor stability assay to evaluate thermostability of POLγA^WT^ (black) and POLγA^A449T^ (blue), in absence (solid line) or presence (dashed line) of POLγB. **C.** Size-exclusion chromatogram of POLγA^WT^ (black line) and POLγA^A449T^ (blue line) in presence of POLγB, to evaluate interaction between POLγA and POLγB. **D.** SDS-PAGE of the selected peak fractions from (C) of POLγA^WT^ and POLγB. **E.** SDS-PAGE of the selected peak fractions from (C) of POLγA^A449T^ and POLγB. Note the brackets highlighting unbound POLγB (free from POLγA^A449T^). **F.** Western blot analysis of steady-state levels of POLγA, LONP1 and POLγB upon siRNA-mediated knockdown of LONP1, POLγB and POLγA, in HeLa cells. ß-actin was used as loading control. **G.** Quantification of POLγA levels upon siRNA-mediated knockdown of LONP1 (F). POLγA levels were normalized to ß-actin and presented as FOLD change from cells treated with control siRNA. Data are presented as mean ± SEM. ***p<0.001; Student’s t-test. (n = 3). **H.** Quantification of POLγA levels upon siRNA-mediated knockdown of POLγB (F). POLγA levels were normalized to ß-actin and presented as fold change from cells treated with control siRNA. Data are presented as mean ± SEM. ***p<0.001; Student’s *t*-test. (n = 3). **I.** Western blot analysis of steady-state levels of POLγA in heart of *Lonp^+/+^* and *Lonp1^−/−^* animals. An anti-LONP1 antibody was used to confirm gene knockout and HSC70 was used as loading control. **J.** Quantification of POLγA levels in heart of *Lonp^+/+^* and *Lonp1^−/−^* animals (I). POLγA levels were normalized to HSC70 and presented as fold change from *Lonp^+/+^*. Data are presented as mean ± SEM. *p<0.05; Student’s *t*-test. (n = 6).

### POLγ^AA449T^ has reduced affinity for POLγB

We hypothesized that the A449T mutation could impair interactions with POLγB and thus destabilize POLγA^A449T^. To address this possibility, we investigated POLγA^A449T^ interactions with POLγB by performing size-exclusion chromatography. At 1:1 molar ratio of POLγA and POLγB (calculated as a dimer), POLγ^WT^ and POLγ migrated as a single peak, corresponding to a stable complex between the two proteins (Figure 6C), as confirmed by SDS-PAGE (Figure 6D). In contrast, POLγA^A449T^ and POLγB showed an additional peak, corresponding to unbound POLγB (Figure 6C and 6E). The resolution of the chromatography cannot separate free POLγA from the POLγ holoenzyme. Thus, the A449T mutation significantly reduces the interaction between POLγA and POLγB subunits. This observation is in agreement with data for human POLγA^A467T^ (Chan et al., 2005).

### POLγB protects POLγA against LONP1 degradation

Next, we investigated if free, partially unfolded POLγA could be a target for protein degradation. In mitochondria, the LONP1 protease degrades misfolded proteins and is linked to regulation of mtDNA copy number (Bezawork-Geleta, Brodie et al., 2015). We therefore decided to investigate if POLγA was a target for LONP1.

We first used siRNA interference against LONP1, POLγA and POLγB in HeLa cells. Interestingly, LONP1 knockdown caused a robust increase in POLγA levels (Figure 6F and 6G), whereas POLγB was unaffected (Figure 6F), supporting the idea that POLγA is a specific target of LONP1 degradation. In agreement with a stabilizing effect of POLγB, knockdown of *Polg2* mRNA also caused a reduction of POLγA levels (Figure 6F and 6H). Both POLγB and POLγA knockdown resulted in an increase of LONP1 (Figure 6F). To further support our findings, we evaluated the steady state levels of PolγA in a mouse model with a tissue-specific Lonp1 knockout (6I and 6J). Notably, PolγA levels were increased in heart samples of *Lonp1*^−/−^ compared to control littermates. Collectively, these results support that LONP1 specifically targets POLγA both in cells and *in vivo*.

To investigate if POLγA is a direct target for LONP1 degradation, we performed a size-exclusion chromatography with recombinant protein to assess if POLγA can form a complex with LONP1. To ensure that PolγA was not degraded by LONP1 during the experiment, we used the mutant LONP1^S855A^, which traps substrates without degrading them (Kereiche, Kovacik et al., 2016). As shown in Figure 7A, we observed a co-elution of LONP ^S855A^ and POLγA, revealing an interaction between these two proteins.

**Figure 7.**
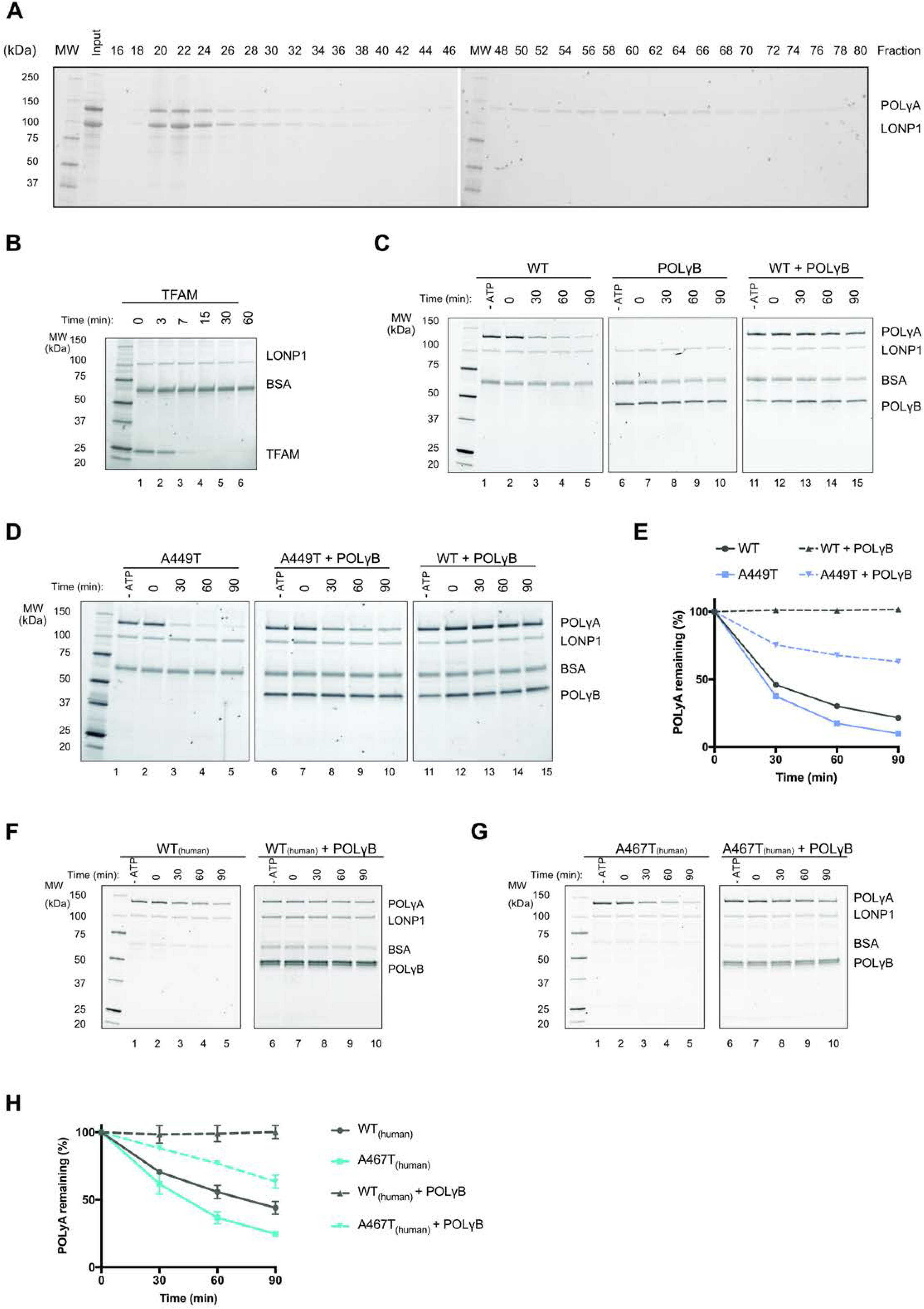
POLγA is a target of LONP1 degradation *in vitro*. **A.** Size-exclusion chromatography of the complex formed by LONP1^S855A^ (Catalytic dead mutant) and WT POLγA. The mixture was incubated for 10min, at 37°C, in the presence of 10 mM MgCl_2_ and 2 mM ATP before loaded on the chromatography. **B.** SDS-PAGE of the LONP1 proteolysis assay of TFAM, over time. The reactions were incubated for the indicated times (top). **C.** SDS-PAGE of the LONP1 proteolysis assay of isolated POLγA^WT^ (left) and POLγB (middle), and POLγA^WT^ complexed with POLγB, over time. The reactions were incubated for the indicated times (top). In the absence of ATP (-ATP control), LONP1 does not exert proteolysis. **D.** SDS-PAGE of the LONP1 proteolysis assay of POLγA^A449T^ in absence (left) or presence (middle) of POLγB, over time. Reactions with WT POLγA + POLγB (right) were added for reference. The reactions were incubated for the indicated times (top). In the absence of ATP (-ATP control), LONP1 does not exert proteolysis. **E.** Quantification of POLγA degradation over time (0-90min) by LONP1, related to (C-D). POLγA^WT^ (black) and POLγA^A449T^ (blue), in absence (solid line) or presence (dashed line) of POLγB. Data are presented as mean ± SD. (n = 3). **F.** SDS-PAGE of the LONP1 proteolysis assay of human POLγA^WT^ in absence (left) or presence (right) of POLγB, over time. The reactions were incubated for the indicated times (top). In the absence of ATP (-ATP control), LONP1 does not exert proteolysis. **G.** SDS-PAGE of the LONP1 proteolysis assay of human POLγA^A467T^ in absence (left) or presence (right) of POLγB, over time. The reactions were incubated for the indicated times (top). In the absence of ATP (-ATP control), LONP1 does not exert proteolysis. **H.** Quantification of human POLγA degradation over time (0-90min) by LONP1, related to (F-G). POLγA^WT^ (grey) and POLγA^A449T^ (blue), in absence (solid line) or presence (dashed line) of POLγB. Data are presented as mean ± SD. (n = 3).

We also monitored LONP1 dependent degradation of POLγA and POLγB *in vitro*. We followed the reactions over time and used another well-characterized LONP1 substrate, TFAM, as a positive control (Lu, Lee et al., 2013, Matsushima, Goto et al., 2010). The TFAM levels were reduced by 50% in about 3 minutes (Figure 7B). We observed no LONP1-dependent degradation of POLγB, confirming that the accessory subunit is not a substrate of LONP1 (Figure 7C, lanes7-10). In contrast, both isolated POLγA^WT^ and POLγA^A449T^ were efficiently degraded, with a 50% reduction in about 20 minutes (Figure 7C, lanes 2-5, 7D, lanes 2-5 and 7E). The slower degradation time compared to TFAM could in part be explained by the size difference between the two substrates, with POLγA being about 6-fold larger. LONP1 is an ATP dependent enzyme, and no degradation of PolγA was therefore observed in the absence of ATP (Figure 7C, lanes 1, 6 and 11).

Next, we examined POLγA in complex with POLγB. Interestingly, the presence of POLγB completely blocked POLγA^wt^ degradation (Figure 7C, lanes 12-15 and Figure 7E). In contrast, POLγB was unable to efficiently block degradation of POLγA^A449T^ and the levels of the mutant protein decreased significantly over the time of the experiment (Figure 7D, compare lanes 7-10 with 12-15 and Figure 7E). We also used the human WT and A467T mutant versions of POLγA, and observed similar results (Figure 7F, 7G and 7H). We conclude that impaired interaction between POLγB and POLγA^A449T^ leads to increased LONP1 dependent degradation of POLγA^A449T^. This observation could explain the lower levels of POLγA^A449T^ observed *in vivo*.

To validate our model, we also analysed two additional POLγA mutations. The mouse version of POLγA^W748S^ (PolγA^W725S^), which also displays reduced interactions with PolγB (Supplementary Figure 6A, 6B and 6C) and human POLγA^D274A^, which has no effect on POLγB interactions (Macao, Uhler et al., 2015). As expected, POLγA^W725S^ but not POLγA^D274A^ was degraded in presence of POLγB (Supplementary Figure 6D, 6E and 6F). Overall, these data provide *in vitro* evidence that POLγB affects POLγA folding and protects the protein from degradation. Our data also suggest that other POLγA mutations affecting the interactions with POLγB or vice versa (e.g. mutations in POLγB affecting the interaction with POLγA) may be subjected to LONP1 degradation.

## Discussion

Mutations in *POLG* are a relatively common cause of a spectrum of mitochondrial disease. The substantial lack of relevant *in vivo* models has hampered our understanding of the pathogenesis of these *POLG*-related disorders. Here we developed a mouse model for human POLG^A467T^ and study the molecular pathogenesis of this common mutation *in vivo.* We complemented this analysis with detailed biochemical characterization of the corresponding events *in vitro*.

The homozygous *Polg^A449T/A449T^* mice displayed a mild decrease of mtDNA in skeletal muscle, impaired treadmill performance and reduced spontaneous movements. The mice also displayed reduced 7S DNA levels in several tissues, as has been previously described in mouse knock-out models for other components of the mitochondrial replication machinery, including POLγB (Di Re, Sembongi et al., 2009); mtSSB (Ruhanen, Borrie et al., 2010); and TWINKLE (Milenkovic, Matic et al., 2013). The obvious explanation for the mild phenotype observed in the *Polg^A449T/A449T^* compared to that of the patients is that compensatory mechanisms may be more efficient in mice than in humans, since the essential biochemical features are qualitatively identical between the two organisms. For instance, one possibility is that in the mouse the stabilization by POLγB is more effective than in humans. In addition, in humans the A467T mutation has been associated to profound depletion of mtDNA in brain and other tissues^10^, while in mice the depletion is very mild, possibly because of more successful replication events, as suggested by reduced 7S DNA levels and increased number of replicating nucleoids in MEFs.

Experiments carried out on *Polg^A449T/A449T^* MEFs showed an increased number of actively replicating nucleoids as well as impaired mtDNA synthesis upon stress-induced mtDNA depletion with EtBr. The observation was in agreement with a reduced capacity of *Polg^A449T/A449T^* mice to recover from the CCl_4_-induced liver necrosis, demonstrating that effective mtDNA replication is necessary for liver regeneration. Similar observations have been made earlier in a knockout mouse model of mitochondrial topoisomerase I (TOP1mt) (Khiati, Baechler et al., 2015). In the *Polg^A449T/A449T^* mice, we observed an accumulation of RIs, as demonstrated both by *in organello* replication assay and 2D-AGE analysis, suggesting a replicative fork stalling phenotype. Notably, we did not observe any deletions in mouse mtDNA. Our analysis of POLγA^A449T^ *in vitro* revealed impaired interactions with POLγB and reduced replication efficiency, which were in agreement with the effects observed *in vivo* and previous *in vitro* analysis of A467T mutation in human POLγA (Chan et al., 2005). Interestingly, a comparison between the mouse POLγA^A449T^ and human POLγA^A467T^ proteins, revealed similar, but more pronounced replication defects for the human polymerase, which may explain why the A467T mutation causes more severe phenotypes, including multiple mtDNA deletions.

As demonstrated here, a reduction of POLγA^A449T^ protein levels also contributes to the pathogenesis of phenotypes observed in the mouse. Using a thermofluor stability assay, we found that POLγA (WT and mutant) was structurally unstable at physiological temperatures, but strongly stabilized in complex with POLγB (Figure 6B). In support to this notion, isolated POLγA was shown to be efficiently degraded by LONP1, a protease that recognizes unfolded mitochondrial proteins, and this degradation was completely blocked in the presence of POLγB. Follow-up experiments *in vivo* supported this observation, since depletion of LONP1 both in a mouse model and in human cells caused an upregulation of POLγA. In addition, depletion of POLγB leads to a concomitant reduction of POLγA levels in HeLa cells. Interestingly, the A449T mutation made the complex between POLγA and POLγB less stable, explaining the decrease of POLγA in our mutant mouse model. This model received further support from our analysis of another mutant form of POLγA, the mouse equivalent of human POLG^W748S^, which also impairs interactions with POLγB.

Our findings clearly indicate that LONP1 degrades POLγA *in vitro* and *in vivo*. However, we cannot exclude that other proteases may also contribute to this process *in vivo*. For instance, ClpXP, another AAA+ protease of the mitochondrial matrix (Goard & Schimmer, 2014) can also degrade free POLγA *in vitro* and the reaction is blocked by addition of POLγB. However, in contrast to LONP1, POLγA levels are not changed in a mouse knock-out model for the ClpXP-protein *(unpublished)*. The rapid degradation of free POLγA could be of physiological relevance, since on its own, the protein displays higher exonuclease activity, which can potentially disturb mtDNA replication (Figure 5C).

In conclusion, we here describe in detail the *in vivo* and *in vitro* features of a common mutation of POLγA, shedding light on the pathogenesis of an intriguing and previously poorly understood condition. Furthermore, we suggest a new mechanism contributing to the pathogenesis of *POLG*-related diseases. Mutations in *POLG* or *POLG2* that cause weaker interactions within the POLγ holoenzyme will lead to LONP1-dependent degradation of POLγA, resulting in protein depletion *in vivo*. We speculate that interventions aimed at increasing POLγA stability, either by increasing interactions with POLγB or by reducing LONP1, may have therapeutic value in affected patients. We believe that the *Polg^A449T/A449T^* mouse model developed here will be a valuable tool for studies of new therapeutic interventions and help clarify the role of mitochondrial proteases in mtDNA maintenance disorders.

## Acknowledgements

Our research is supported by the Telethon Foundation (grant GGP19007 to MZ), NRJ-Institute de France Grant (to M.Z.), Fondazione Onlus Luigi Comini (to MZ and CV); and core grants from the Medical Research Council (Grant MC_UU_00015/5, MC_UU_00015/4 and MC_UU_00015/7). PSP, CPH and DHH are supported by the Marie Sklodowska-Curie ITN-REMIX grant (Grant 721757), the Swedish Research Council (MF); Swedish Cancer Foundation (MF); European Research Council (MF); the Knut and Alice Wallenbergs Foundation (MF). LT is supported by a MRC-funded graduate student fellowship. SAD is supported by EMBO Installation Grant IG4149, TUBITAK 119C022 and Bogazici University SUP-15501.

## Authors’ contribution

MZ, LAB, MF, BM and CV designed the study, PP and SAD performed the *in vivo* experiments, PP and CPH performed the biochemical and molecular analysis, PP and AR performed the 2D-AGE experiments, RC performed the histological and histochemical analysis; AM, DHH performed the *in vivo* and *in vitro* experiments with LONP1; SL and LT performed the experiments on MEFs; PFS contributed to design the mouse model and to perform the BNGE analysis; AT supervised DHH for the LONP1 mouse model; MM supervised PP during the last part of the project; JP supervised LT. All the authors contributed to writing the manuscript.

## Methods

### Generation of *Polg^A449T/A449T^* mice

*Polg^A449T/A449T^* mice were generated by a double-nickase CRISPR/Cas9 D10A mediated gene editing of mouse *Polg* gene in exon 7 (c.1345G>A / p.A449T). For a detailed representation see (Supplementary Figure 1A). The selected sgRNAs (Table 1) were cloned into plasmid pSpCas9(BB)-PX330 (Addgene #42230), using the *BbsI* site. The resulting constructs were used as a template to amplify by PCR the gRNA (spacer + scaffold) preceded by a T3 promoter to allow subsequent *in vitro* transcription. The *in vitro* transcription was carried using the MEGAscript T3 Transcription Kit (Life Technologies). The same kit was used to produce Cas9 D10A mRNA using as template the plasmid pCAG-T3-hCasD10A-pA (Addgene #51638). The 140bp ssDNA homology direct repair (HDR) donor (Table 1) was acquired from IDT. Cas9 D10A mRNA, gRNAs and HDR donor were microinjected into fertilized FVB/NJ one-cell embryos (Core Facility for Conditional Mutagenesis, Milan). Genotyping of *Polg^A449T/A449T^* mice was performed by PCR (primers Polg_A449T_Fw + Polg_A449T_Rv, Table 1), followed by a restriction digestion with *PvuII*. WT allele produces a fragment of 769bp, which is cleaved in the A449T allele producing two fragments of 490bp + 279bp (Supplementary Figure 1B). The PCR is carried using GoTaq DNA polymerase (Promega, UK) and the following PCR conditions: 95 °C for 30 s, 63.7 °C for 30 s, and 72 °C for 1 min, 35 cycles.

**Table 1.**
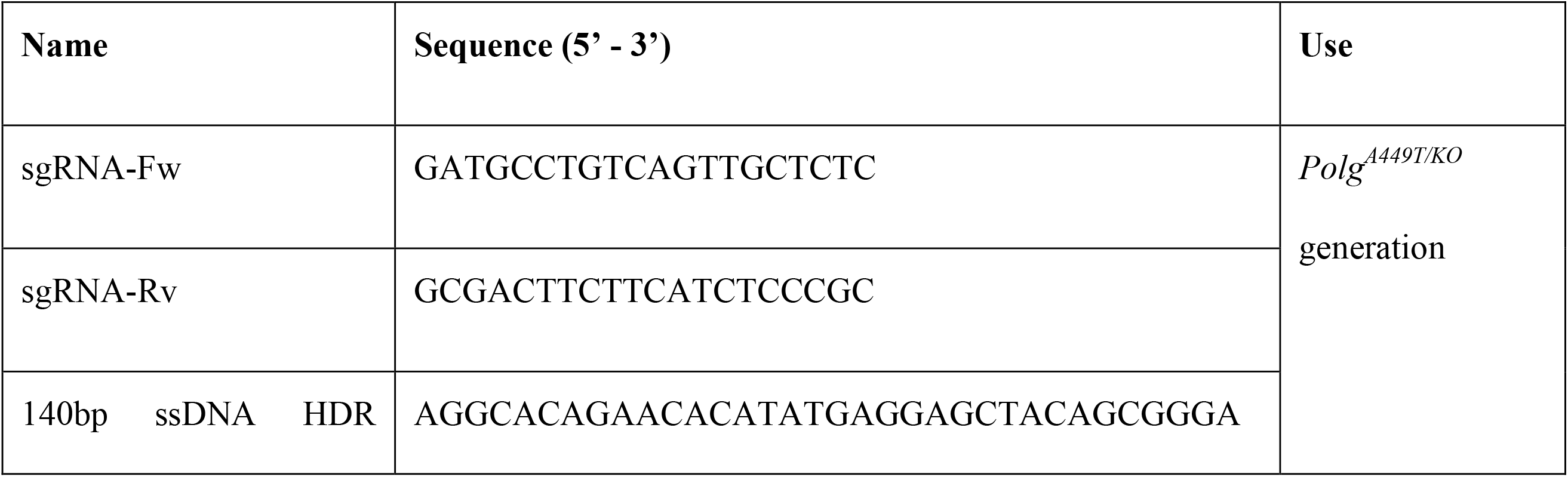
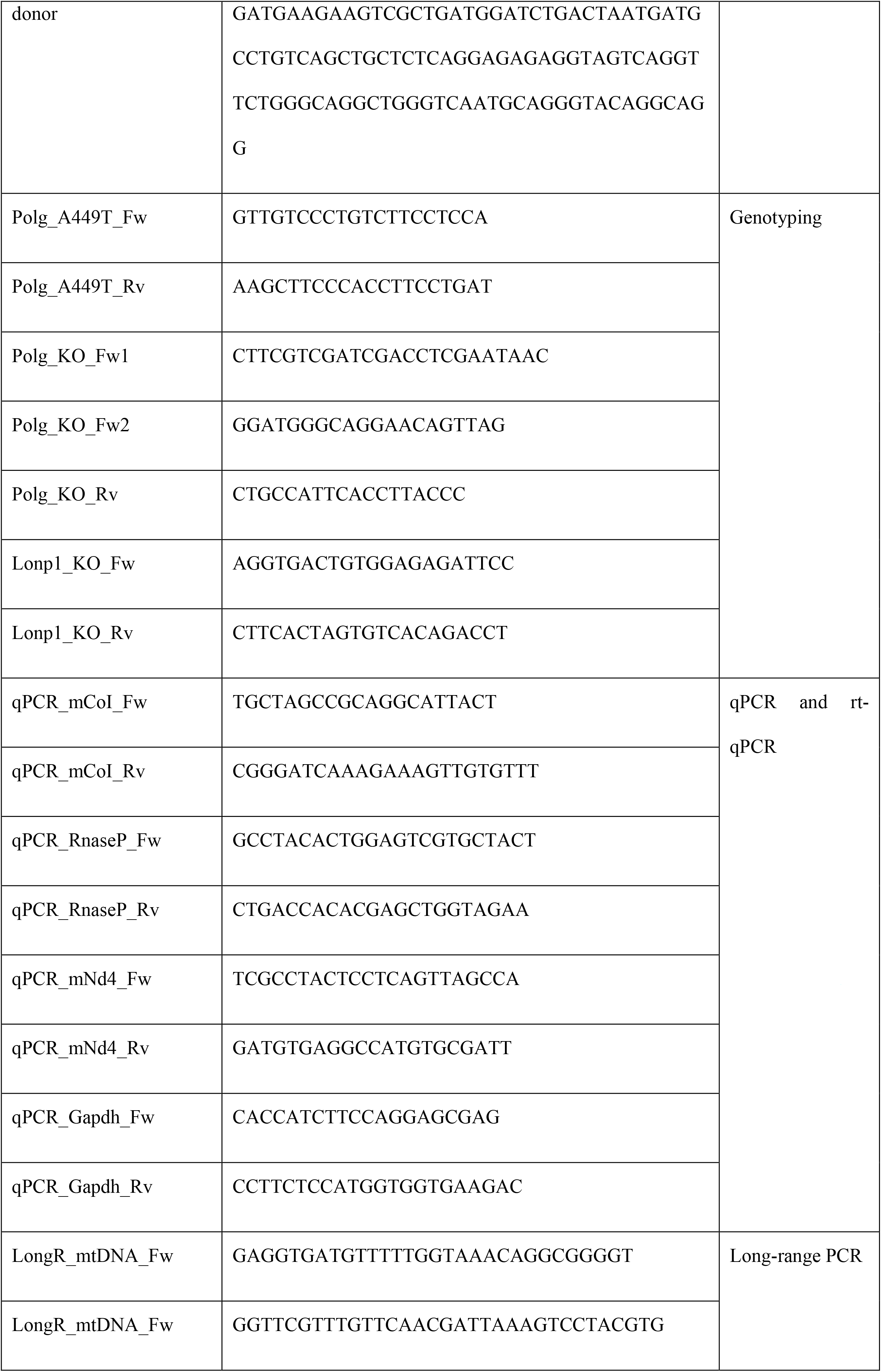
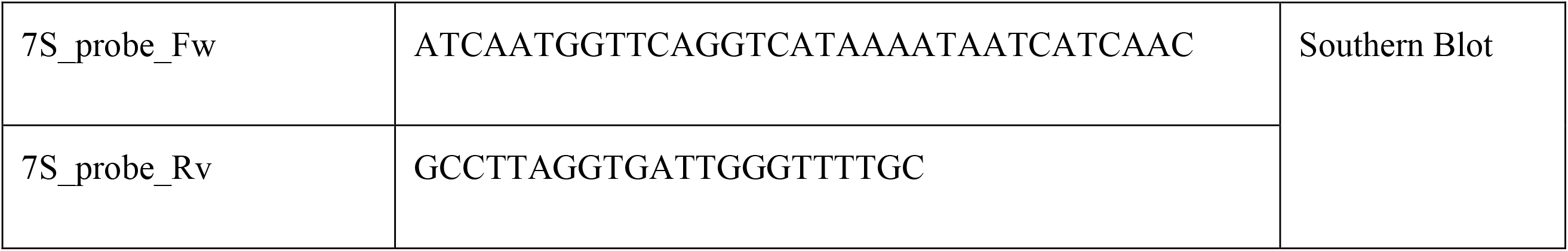
Oligonucleotides list.

### Animal work

All animal experiments were carried out in accordance with the UK Animals (Scientific Procedures) Act 1986 (PPL: P6C20975A) and EU Directive 2010/63/EU. The mice were kept on FVB/NJ background, and wild-type littermates were used as controls. The animals were maintained in a temperature- and humidity-controlled animal care facility with a 12-h light/12-h dark cycle and free access to water and food, and they were monitored weekly to examine body condition, weight, and general health. All mice were sacrificed by cervical dislocation at 3 months of age for subsequent analysis.

*Lonp1* gene targeting (*Lonp1^+/tm1a(EUCOMM)Hmgu/Ieg^,* project number HEPD0936_3_B11) was carried out as part of the The European Conditional Mouse Mutagenesis Program (EUCOMM), on the C57BL/6NTac genetic background. We generated the heart and skeletal muscle specific *Lonp1* knockout mice by mating *Lonp1^fl/fl^* animals with transgenic mice expressing cre recombinase under the control of muscle creatine kinase promoter (*Ckmm-cre*) (Larsson, Wang et al., 1998), after removal of a gene-trap DNA cassette. Experiments were performed on 12-week-old mice. The genotyping primers used are on Table 1 (Lonp1_KO_Fw + Lonp1_KO_Rv).

All experiments on *Lonp1^fl/fl^; Ckmm-Cre* animals were approved and permitted by the Animal Ethics Committee of North-Rhein Westphalia (Landesamt für Natur, Umwelt und Verbraucherschutz Nordrhein-Westfalen; LANUV) following the German and European Union regulations.

### Treadmill

A standard treadmill apparatus (Panlab) was used to measure motor endurance according to the number of falls in the motivational air puff during a gradually accelerating program with speed initially at 6.5 m/min and increasing by 0.5 m/min every 3 min. The test was terminated by exhaustion, defined as >10 air puffs activations/min.

### Comprehensive laboratory animal monitoring system (CLAMS)

Mice were individually placed in the CLAMS™ system of metabolic cages and monitored over a 48-h period. Data were collected every 10-min. The parameters analyzed were: ambulatory and rear movements, VO2 (volume of oxygen consumed, ml/kg/h), VCO2 (volume of carbon dioxide produced, ml/kg/h), RER (respiratory exchange ratio) and heat (kcal/h).

### Pharmacological treatments

In VPA-treated mice, VPA (Sigma) was administrated by daily oral gavage (300mg/kg in water) or added to a standard diet at 1.5% (1.5g-VPA/ 1kg-Food) and administered for 60 days, starting at 8 weeks of age.

In CCl_4_ experiments, mice received a single IP injection of CCl_4_ (1 mL/kg body weight diluted 1/10 in olive oil (Sigma). Mice were sacrificed after 2 or 4 days. For histology analysis, Livers samples were fixed in 10% formalin and embedded in paraffin. Sections of 4-μm were stained with H&E according to standard protocols. The quantification of necrotic areas was done with ImageJ by dividing the necrotic areas around the central veins by total area of the section. Five different regions of the slide were analyzed and average value obtained.

### DNA and RNA extraction

Genomic DNA was extracted by resuspending samples in lysis buffer (0.5% sodium dodecyl sulfate (SDS); 0.1M NaCl; 50mM Tris-HCl, pH=8; 2.5mM EDTA). Samples were incubated overnight at 65 °C after adding Proteinase K (final concentration: 20ng/μL). Next, samples were purified with 1 volume chloroform + 0.6M potassium acetate and supernatant was ethanol precipitated. Final DNA was eluted in water.

Total RNA was extracted from the indicated tissues using the TRIzol Reagent (Thermofisher) following the manufacture protocol.

### Real-time quantitative PCR

For mtDNA relative quantification, SYBR Green real-time qPCR was performed using primers specific to a mouse mtDNA region in the COI gene. Primers specific to RNaseP, a single copy gene taken as a nuclear gene reference. All primers are listed in Table 1. Approximately 25 ng of DNA was used per reaction.

For the quantification of mRNA levels, cDNA was retrotranscribed from total RNA extracted using the Omniscript RT kit (Qiagen). For mitochondrial transcripts *CoI* and *Nd4*, specific primers (Table 1) were used as described above with SYBR Green chemistry. Expression was calculated using the ΔΔCt analysis using *Gapdh* as reference.

Specific Gene Expression TaqMan assays (Invitrogen) were used for *Polg* and *Polg2*. Expression was calculated using the ΔΔCt analysis using *B2m* as reference.

### Long-range PCR

MtDNA was amplified from 50 ng of total DNA with the primers (LongR_mtDNA_Fw and LongR_mtDNA_Rv, Table 1) using PrimeSTAR GXL DNA polymerase (TAKARA, Japan) and following PCR conditions: 98 °C for 10 s, 68 °C for 13 min, 35 cycles.

### Cell cultures

*Polg^A449T/A449T^* and control mouse embryo fibroblasts (MEFs) were prepared from individual E12.5 embryos and were cultured in complete DMEM with high glucose and 10% fetal bovine serum. MEFs were seeded in six-well plates at 20% confluence. Cells were incubated with or without 100ng/mL EtBr for 5 days and DNA samples were collected every 24h. At day 5, cells new medium without EtBr was added and cells were allowed to recover for an additional 8 days. Again, DNA samples were collected every 24h. MtDNA quantification was performed as described above.

HeLa cells were grown at 37°C, 5% CO2 in Dulbecco’s Modified Eagle Medium (DMEM; 4.5 g/L glucose, 2 mM glutamine, 110 mg/ml sodium pyruvate) supplemented with 10% fetal bovine serum (FBS) and 5% penicillin/streptomycin. For siRNA transfections, 0.3 × 10^6^ HeLa cells were reverse transfected with 5 nM of siRNA using Lipofectamine RNAiMAX. siRNAs used in the study are: (i) LONP1 (5′-GGUGCUGUUCAUCUGCACGtt-3′), (ii) PolgB (5′-CGGUGCCUUGGAACACUAUtt-3′), (iii) PolgA (5′-CCCAUUGGACAUCCAGAUGtt-3′). After three days, cells were harvested, washed with PBS and used for Western Blotting as described above.

### Immunofluorescence analysis and confocal imaging

Immunofluorescence were performed as previously described (Nagashima, Tabara et al., 2020). Briefly, cells seeded in 24-well plate were fixed in 5% paraformaldehyde (PFA) in PBS at 37°C for 15 minutes (min) and incubated with 50 mM ammonium chloride in PBS for 10 min at room temperature (RT). After three washes in PBS, cells were permeabilized using 0.1% Triton X-100 in PBS for 10 min, washed 3 times with PBS, and then blocked in 10% FBS in PBS for 20 min at RT. Cells were then incubated with indicated primary antibodies for 2 hours in 5% FBS/PBS, washed in 5 % FBS in PBS and incubated with secondary Alexa Fluor conjugated antibodies in 5% FBS/PBS for 1 hour at RT. We used the following antibodies: TOM20 (1:1000) was from Abcam (ab232589), DNA (1:1500) was from Millipore (CBL186), Goat Anti-rabbit Alexa Fluor 594 (1:1000) was from Invitrogen (A-11012), Donkey anti-mouse Alexa Fluor 488 (1:1000) was from Invitrogen (A-21202). EdU incorporation was detected using Invitrogen Click-iT EdU AlexaFluor 647 (Invitrogen, C10340) labelling kit according to manufacturer’s instructions. Coverslips were mounted onto sides using Dako fluorescence mounting medium (Dako). Images were then acquired as 7 stacks of 0.2 μm each, using a 100X objective lense (NA1.4) on a Nikon Eclipse TiE inverted microscope using an Andor Dragonfly 500 confocal spinning disk system, equipped with a Zyla 4.2 PLUS sCMOS camera, exciting with 488 nm, 594 nm or 633 nm lasers, and coupled with Fusion software (Andor). For quantification of EdU or mtDNA number, max projection images were processed once with the “smooth” function in Fiji and nucleus was removed. Images were then manually thresholded, ‘smoothed’ and number of particles were obtained using the “Analyze particles” plugin in Fiji with a minimum area of 0.1 μm^2^. The representative images in figure 3 were processed once with the “smooth” function in Fiji.

### Biochemical analysis of MRC complexes

Liver and muscle samples stored in liquid nitrogen were homogenized in 10mM of potassium phosphate buffer (pH=7.4), and the spectrophotometric activity of respiratory chain complexes I, II, III and IV, as well as citrate synthase, was measured as described (Bugiani, Invernizzi et al., 2004).

### BNGE and in-gel activity

For blue native gel electrophoresis (BNGE) analysis, skeletal muscle and liver mitochondria were isolated as previously described (Fernandez-Vizarra, Lopez-Perez et al., 2002). Samples were resuspended in 1.5 M aminocaproic acid, 50 mM Bis-Tris/HCl (pH 7) and 4 mg of dodecyl maltoside/mg of protein, and incubated for 5 min on ice before centrifuging at 20,000 × g at 4°C. 5% Coomassie G250 was added to the supernatant. 100 μg was separated by 4%–12% gradient BNGE and further subjected to a Complex I in-gel activity (IGA), as previously described (Calvaruso, Smeitink et al., 2008). To allow for cI activity to appear, gels were incubated between 1.5 and 24 h in cI-IGA reaction buffer.

### COX/SDH histochemical analysis

For histochemical analysis, skeletal muscle samples were frozen in isopentane pre-cooled in liquid nitrogen. Eight-micrometer-thick sections were stained for COX, SDH and combined COX/SDH activity as described (Sciacco & Bonilla, 1996).

### Southern Blot

Three micrograms of total DNA isolated from each tissue were restricted using the restriction enzyme *BlpI* according to manufacturer’s instructions (New England Biolabs). Products were separated on 0.8% agarose gels (Invitrogen Ultrapure) and dry-blotted overnight onto nylon membrane (GE Magnaprobe). Membranes were hybridized with radiolabeled probes overnight at 65°C in 0.25 M phosphate buffer (pH 7.6) and 7% SDS, then washed for 3 × 20 min in 1× SSC and 0.1% SDS and imaged using a phosphorimager (GE Healthcare) and scanned using an Amersham Typhoon 5 scanner. For primer sequences used for producing probes, see Table 1.

### In Organello Replication

Labeling of mtDNA in isolated organelles was performed as previously described (Reyes, Kazak et al., 2013).

Briefly, isolated liver was minced and homogenized in 4 ml/g of tissue in Sucrose-Tris-EDTA (STE)-buffer [320 mM sucrose, 10 mM Tris–HCl (pH 7.4), 1 mM EDTA and 1 mg/mL essentially fatty acid–free bovine serum albumin (BSA)] using a manual tight-fitting teflon pestle. Resulting mitochondria were washed once in STE-buffer, pelleted and equilibrated in incubation buffer [10 mM Tris–HCl (pH 8.0), sucrose and glucose 20 mM each, 65 mM D-sorbitol, 100 mM KCl, 10 mM K 2 HPO 4, 50 μM EDTA, 1 mg/mL BSA, 1 mM ADP, MgCl 2, glutamate and malate 5 mM each]. In organello labeling was performed for 5; 15; 30; 60 and 90 minutes, at 37°C with rotation, using 1 mg/mL mitochondria in incubation buffer supplemented with dCTP, dGTP and dTTP (50 μM each) and [α-^32^P]-dATP (Hartmann, 3000 Ci/mmol) at 6.6 nM. At the end, DNA was extracted by solubilizing mitochondria with 1% sodium N-lauroylsarcosinate, followed by 100 μg/mL Proteinase K on ice for 30 min and phenol-chloroform extraction. Gel electrophoresis, southern blotting and hybridization were carried as described above.

### 2D-AGE

For two-dimensional gels, DNA was extracted from fresh liver-isolated mitochondria purified by sucrose gradient followed by phenol-chloroform extraction. Five micrograms of the resulting mtDNA were restricted digested with *BclI* according to manufacturer’s instructions (New England Biolabs). For first dimension, products were separated on 0.4% agarose gels (Invitrogen Ultrapure) without ethidium bromide. Then each lane was excised and rotated 90° anticlockwise for second dimension electrophoresis by casting around the gel slices 1% agarose with 500 ng/mL ethidium bromide. After electrophoresis, southern blotting and hybridization were carried as described above.

### Western Blot and Antibodies

Mouse tissues were homogenized in RIPA buffer [150 mM sodium chloride, 1.0% NP-40, 0.5% sodium deoxycholate, 0.1% SDS (sodium dodecyl sulfate), 50 mM Tris, pH 8.0] in the presence of protease inhibitors (cOmplete™ Protease Inhibitor Cocktail, Sigma). Protein concentration was determined by the Lowry method. Aliquots, 30 μg each, were run through a 12% SDS-PAGE and electroblotted onto a polyvinylidene fluoride (PVDF) membrane, which was then immunodecorated with different primary antibodies: anti-POLγA (1:500) was from Santa Cruz Biotechnology (sc-5931), anti-POLγB (1:1000) was from LSBio (LS-C334882), anti-GAPDH (1:3000) was from Abcam (ab53098), anti-LONP1 (1:1000) was from Proteintech (15440-1-AP), anti-HSC70 (1:1000) was from Santa Cruz Biotechnology (sc-7298). Secondary antibodies were from Promega (catalog nos. W4011 [rabbit], W4021 [mouse] and V8051 [goat]). HeLa cells were lysed in lysis buffer (0.125 M Tris HCl, pH. 6.8., 4% SDS and 500mM NaCl). Whole cell lysates were quantified and 50 μg were resolved in 4-20% SDS-PAGE and transferred onto nitrocellulose membranes (GE healthcare). The membranes were then incubated with the primary antibodies: anti-PolgA (1:1000) was from Abcam (ab128899), anti-PolgB (1:500) was home-made, anti-LONP1 (1:1000) was from Abcam (cat ab103809), anti-β actin (1:10000) was from Abcam (ab6276).

### Production of LONP1, POLγA and POLγB, expression and purification

Wild-type LONP1 gene lacking the mitochondrial targeting sequence (aa 1-67) was cloned into a pNic28-BSA4 vector with a cleavable 6xHisTag in the N-terminus. Rosetta™(DE3) pLysS competent cells (Novagen) were transformed with the plasmid and grown in Terrific Broth media with 50 mg/l Ampicillin and 34 mg/l Chloramphenicol at 37°C until OD_600_ = 3. Protein expression was induced with 1 mM IPTG at 16°C for 4 h.

Cells were harvested by centrifugation, frozen in liquid nitrogen, thawed and lyzed at 4°C in lysis buffer (25 mM Tris-HCl pH 8.0, 10 mM β-mercaptoethanol). The suspension was homogenized using an Ultra-Turrax T3 homogenizer (IKA) and centrifuged at 20000 × g for 45 min in a JA-25.50 rotor (Beckman Coulter). The supernatant was loaded onto His-Select Nickel Affinity Gel (Sigma-Aldrich) equilibrated with buffer A (25 mM Tris–HCl, pH 8.0, 0.4 M NaCl, 10% glycerol and 10 mM β-mercaptoethanol). The protein was eluted with buffer A containing 250 mM imidazole. Removal of the 6xHis tag was achieved by overnight-dialysis in presence of ≈ 0.5 mg TEV in buffer A. An additional Nickel purification step performed in order to get rid of uncut His tagged protein and TEV. The protein was subsequently purified over a 5 ml HiTrap Heparin HP column (GE Healthcare) and a 1 ml HiTrap Q HP column (GE Healthcare), both equilibrated in buffer B (25 mM Tris-HCl pH 8.0, 10% glycerol and 1 mM DTT) containing 0.2 M NaCl, followed by elution driven by a linear gradient (50 and 10 ml respectively) of buffer B containing 1.2 M NaCl (0.2-1.2 M NaCl). Protein purity was checked on a precast 4-20% gradient SDS-PAGE gel (BioRad, 567-8094) and pure fractions were aliquoted and stored at −80°C.

POLγA and POLγB were expressed and purified as described previously (Farge, Pham et al., 2007), with the following modifications: For POLγA, an additional step of purification with 1 ml HiTrap SP HP column was added after the HiTrap Q HP column purification. The column was equilibrated with buffer B containing 0.1 M NaCl and eluted with a linear gradient (10 ml) of buffer B containing 1.2 M NaCl (0.1-1.2 M NaCl). For POLγB, an additional step of purification with a 1 ml HiTrap Talon column (GE Healthcare) was used in between HiTrap Heparin HP (GE Healthcare) and HiTrap SP HP (GE Healthcare). This column was equilibrated with buffer C (25 mM Hepes pH 6.8, 10% glycerol, 0.4 M NaCl, 1 mM β-mercaptoethanol) containing 5 mM imidazole and elution driven by a linear gradient (10 ml) of buffer C containing 150 mM imidazole (5 mM-150 mM imidazole).

For the generation of mutant versions of POLγA, QuikChange Lightning Site-Directed Mutagenesis Kit (Agilent, #210519) was used according to manufacturer’s indications.

### Electrophoresis mobility shift assay (EMSA)

DNA binding affinity of POLγA and POLγA-B2 to a primer-template was assayed using a 36-nt oligonucleotide [5′-TTTTTTTTTTATCCGGGCTCCTCTAGACTCGACCGC-3′] annealed to a ^32^P 5′-labeled 21-nt complementary oligonucleotide (5′-GCGGTCGAGTCTAGAGGAGCC-3′). This produces a primed-template with a 15 bases single-stranded 5′-tail. Reactions were carried out in 15 μl volumes containing 10 fmol DNA template, 20 mM Tris-HCl [pH 7.8], 1 mM DTT, 0.1 mg/ml bovine serum albumin, 10 mM MgCl_2_, 10% glycerol, 2 mM ATP, 0.3 mM ddGTP and 3 mM dCTP. POLγA and POLγB were added as indicated in the figures and reactions were incubated at RT for 10 min before separation on a 6% Native PAGE gel in 0.5 × TBE for 35min at 180V. Bands were visualized by autoradiography.

For K_d_ analysis, band intensities representing unbound and bound DNA were quantified using Multi Gauge V3.0 software (Fujifilm Life Sciences). The fraction of bound DNA was determined from the background-subtracted signal intensities using the expression: bound/(bound+unbound). The fraction of DNA bound in each reaction was plotted versus the concentration of POLγA or POLγA-B2. Data were fit using the “one site – specific binding” algorithm in Prism 8 (Graphpad Software) to obtain values for K_d_.

### Coupled exonuclease-polymerase assay

DNA polymerization and 3′-5′ exonuclease activity were assayed using the same primer-template as described above for EMSA. The reaction mixture contained 10 fmol of the DNA template, 25 mM Tris-HCl [pH 7.8], 10% glycerol, 1 mM DTT, 10 mM MgCl_2_, 100 μg/ml BSA, 60 fmol of POLγA, 120 fmol of POLγB and the indicated concentrations of the four dNTPs. The reaction was incubated at 37°C for 15 min and stopped by the addition of 10 μl of TBE-UREA-sample buffer (BioRad). The samples were analysed on a 15% denaturing polyacrylamide gel in 1 × TBE buffer.

### DNA synthesis on ssDNA template

A ^32^P 5′-labeled 70-mer oligonucleotide [5′-42(T)-ATCTCAGCGATCTGTCTATTTCGTTCAT-3′] was hybridized to a single-stranded pBluescript SK(+). The template formed consists of a 42 nt single-stranded 5′-tail and a 28 bp duplex region. Reactions were carried out in 20 μl volumes containing 10 fmol template DNA, 25 mM Tris–HCl (pH 7.8), 1 mM DTT, 10 mM MgCl_2_, 0.1 mg/ml BSA, 100 μM dATP, 100 of the four dNTPs, 2.5 pmol mtSSB, 150 fmol POLγA and 300 fmol POLγB. Reactions were incubated at 37°C for the indicated times and stopped by the addition of 6 μl of stop buffer (90 mM EDTA, 6% SDS, 30% glycerol, 0.25% bromophenol blue and 0.25% xylene cyanol) and separated on a 0.9% agarose gel at 130V in 1× TBE for 4h.

### Rolling circle *in vitro* replication assay

A ^32^P 5′-labeled 70-mer oligonucleotide [5′-42(T)-ATCTCAGCGATCTGTCTATTTCGTTCAT-3′] was hybridized to a single-stranded pBluescript SK(+) followed by one cycle of polymerization using KOD polymerase (Novagen) to produce a ∼3-kb double-stranded template with a preformed replication fork. Reactions of 20μl were carried out containing 10 fmol template DNA, 25 mM Tris–HCl (pH 7.8), 1 mM DTT, 10 mM MgCl2, 0.1 mg/ml BSA, 4 mM ATP, 100 μM dATP, 100 μM dTTP, 100 μM dGTP, 10 μM dCTP, 2 μCi [α-^32^P] dCTP, 2pmol mtSSB, 200 fmol TWINKLE, 200 fmol POLγA and 500 fmol POLγB (or as indicated in the figure). Reactions were incubated at 37°C for 60 min (or as indicated in the figure) and stopped with 6 μL alkaline stop buffer (18% [wt/vol] Ficoll, 300 mM NaOH, 60 mM EDTA [pH8], 0.15% [wt/vol] Bromocresol green, and 0.35% [wt/vol] xylene cyanol). Products were run in 0.8% alkaline agarose gels and visualized by autoradiography.

Incorporation of [α-^32^P]-dCTP was measured by spotting 5 μl aliquots of the reaction mixture (after the indicated time points at 37°C) on Hybond N+ membrane strips (GE Healthcare Lifesciences). The membranes were washed (3 × with 2 × SSC and 1 × with 95% EtOH) and the remaining activity was quantified using Multi Gauge V3.0 software (Fujifilm Life Sciences). A dilution series of known specific activity of [α-^32^P]-dCTP was used as a standard.

### Thermofluor assay

The fluorescent dye Sypro Orange (Invitrogen) was used to monitor the temperature-induced unfolding of wild-type and mutant POLγA. Wild type and mutant proteins were set up in 96-well PCR plates at a final concentration of 1.6 μM protein and 5× dye in assay buffer (50 mM Tris–HCl pH 7.8, 10 mM DTT, 50 mM MgCl2 and 5 mM ATP). Differential scanning fluorimetry was performed in a C1000 Thermal Cycler using the CFX96 real time software (BioRad). Scans were recorded using the HEX emission filter (560–580 nm) between 4 and 95°C in 0.5°C increments with a 5 s equilibration time. The melting temperature (Tm) was determined from the first derivative of a plot of fluorescence intensity versus temperature (Matulis, Kranz et al., 2005). The standard error was calculated from 3 independent measurements.

### LONP1 proteolysis assay

Protease activity of purified LONP1 on POLγA was measured in a 15 μl reaction volumes containing 0.5 μg of LONP1 wild-type and 0.55 μg of POLγA (in presence or absence of 0.22 ug of PolγB). When having both POLγA and POLγB in the same reaction, a preincubation in ice for 10 min is made before adding LONP1 to the reaction. Samples were incubated at 37°C for 0-90 min in a buffer containing 50 mM Tris-HCl pH 8.0, 10 mM MgCl_2_, 0.1 mg/ml BSA, 2 mM ATP and 1 mM DTT and the reactions were stopped by addition of Laemli sample buffer (BioRad). Samples were run on precast 4-20% gradient SDS-PAGE gels (BioRad, 567-8094) and visualized using ImageLab™ (BioRad) in order to detect proteolytic activity on POLγA. Band intensities were measured with ImageLab™ (BioRad) and calculations were made in order to provide % remaining POLγA-values. Reactions and calculations were made in triplicate and SD was calculated.

### Gel filtration analysis

Complex formation between POLγA and POLγB was tested by size-exclusion chromatography using a Superose 6 Increase 10/300 column (GE Healthcare) connected to an ÄKTA Purifier (GE Healthcare). The column was equilibrated in buffer D (25 mM Tris–HCl, pH 7.8, 10% glycerol, 1 mM DTT, 0.5 M NaCl, 10 mM MgCl_2_). Equal amounts (1 nmoles) of PolγA and PolγB were pre-incubated in buffer D for 10 min on ice before injection. Samples (200 μl) were injected onto the column through a 200 μl loop and run at 1ml/min. Fractions of 250 μl were collected and analyzed on a precast 4-20% gradient SDS-PAGE gel and visualized using ImageLab™ (BioRad). A size calibration curve was previously prepared using thyroglobulin (670 kDa), c-globulin (158 kDa), ovalbumin (44 kDa), myoglobin (17 kDa) and vitamin B12 (1.35 kDa) according to the manufacturer’s instructions (BioRad, 151-1901).

To analyze the LONP1-POLγA interaction, a home-made gel filtration column (0,5 cm × 30 cm) was prepared using Bio-Gel agarose with a bead size of 75-150 μm (BioRad, 151-0440) and calibrated using a gel filtration standard (BioRad, 151-1901). Preincubation of the proteolytic mutant of LONP1 with PolγA in the presence of 2 mM ATP and 10 mM MgCl_2_ at 37°C for 10 min allowed the formation of the complex that was later injected into the column and eluted in 1 CV of buffer D. Fractions of 200 μl were collected and analyzed in a precast 4-20% gradient SDS-PAGE gel and visualized using ImageLab™ (BioRad).

### Statistical analysis

All numerical data are expressed as mean ± SEM unless otherwise stated. A Student’s *t*-test was used to assess statistical significance (see figure legends for details) in two groups comparisons. Differences were considered statistically significant for p <0.05. Animals were randomized in treated and untreated groups. No blinding to the operator was used.

